# Problem-solving without a cortex: inferior lobe drives goal-directed object manipulation in cichlid fish

**DOI:** 10.1101/2025.06.10.658878

**Authors:** Pierre Estienne, Gwenaël Pagé, George Poon, Benoit Larrat, Théo Durier, Ambre Dufresne, Luisa Ciobanu, Kei Yamamoto

## Abstract

Goal-directed object manipulation and problem-solving, which are necessary to evolve tool use behaviors, have long been linked to the expansion of the telencephalon in mammals and birds. Here, we show that goal-directed object manipulation in cichlid fish is driven by a non-telencephalic brain structure, the inferior lobe. Using manganese-enhanced MRI (MEMRI) on an ultra-high field 17.2 Tesla MRI system, we show that the inferior lobe is activated during a puzzle-box opening task. Furthermore, magnetic resonance (MR)-guided High Intensity Focused Ultrasound (HIFU) lesions profoundly impair fine motor coordination during this task without affecting general locomotion or motivation. These results reveal that cortex-like cognitive functions can arise from non-telencephalic brain structures in teleosts. With no homolog in tetrapods, the inferior lobe is a critical hub for flexible behavior in teleost fish. Our findings highlight the existence of alternative neural architectures for the emergence of complex cognition.

## INTRODUCTION

The ability to use tools, an example of higher-order cognitive function, requires innovating to find a solution to a novel situation^1^, and has evolved multiple times independently in mammals and birds^2,3^. Tool-using abilities have been linked to the enlargement of the pallium (the dorsal part of the telencephalon, which contains the cerebral cortex in mammals) in mammals and birds^4^, with primates, crows and parrots being the most proficient^5^ tool users. Tool-using behaviors require multiple cognitive modules including problem-solving, long duration working memory and flexibility^5^.

A shared feature of all species capable of problem-solving is the presence of fine motor control, allowing for the manipulation of objects with the hand, paw, foot, beak or tongue, a prerequisite to execute such goal-directed object manipulation tasks^6,7^. We observe differing degrees of ability for fine motor control between species, which seems to correlate with the degree of problem-solving abilities^5^. For instance, in comparison to primates, carnivores display problem-solving behaviors that seem less flexible^8,9^. This might be due to the lack of efficient grasping appendages in carnivores, as the paw of a dog is less precise than the hand of a macaque^10^. Besides mammals and birds, few species display such manipulation abilities. In cephalopods, octopus can produce extremely precise arm movements to grasp objects and have been shown to possess problem-solving abilities, solving a push-pull puzzle and opening jars with screwed down lids^11–13^.

In teleost fish, problem-solving with object manipulation may be made difficult by the lack of prehensile limbs. Nonetheless, several species of the wrasse (*Labridae*) family are known to use tools^14–18^, requiring great coordination and manipulation abilities using the mouth. Cichlids (*Cichlidae*) have similar manipulation abilities displayed in their building of nests and play behaviors^5,19,20^. However, no study has so far investigated the neural substrate of higher-order cognitive functions in teleosts. As teleost brains are markedly different from that of tetrapods, with many structures such as the telencephalon, hypothalamus, and sensory pathways^21–26^ being much less conserved than previously thought^25,27,28^, the question of the neural correlates of these cognitive functions in teleosts is of great interest to understand the evolution of cognition in vertebrates.

In a previous study, we identified a brain region that was massively enlarged and abundantly connected with the pallium in tool-using wrasses and cichlids, but not in other teleosts^29^. This structure, the inferior lobe, has long been erroneously considered to be part of the hypothalamus, but most of its cells are of mesencephalic origin^21^. As there is no homolog for the inferior lobe in tetrapods, studies about its function remain scarce. It has been suggested that the inferior lobe, in association with the pallium, might be involved in higher-order cognitive functions^29–31^. Indeed, another recent publication highlighted the role of the inferior lobe in a visual learning task in cichlids, a function traditionally associated with the pallium in amniotes^32^. In our past study, we hypothesized that given its many sensory afferents (visual^31,33–35^, somatosensory^31^, and gustatory^30,34,36,37^) and its connectivity with the cerebellum^34,38^, the inferior lobe might also play a role in the fine sensory-motor coordination of the mouth required for delicate object manipulation as observed in tool-use or nest construction behaviors^21,29^.

To verify this hypothesis, we designed a behavioral task requiring goal-directed object manipulation in the convict cichlid (*Amatitlania nigrofasciata*). Using Manganese-Enhanced MRI (MEMRI), a form of functional imaging, we identified increased neuronal activity in the inferior lobe during this behavior. We further confirmed the role of the inferior lobe in goal-directed object manipulation by lesioning it using magnetic resonance (MR)-guided High Intensity Focused Ultrasound (HIFU), which produced significant impairments in task completion. Our results demonstrate that goal-directed object manipulation tasks that involve the telencephalon in mammals^39,40^ involve non-telencephalic structures in teleosts. This further supports the idea that different brain regions can be recruited in distant clades during evolution to perform similar behaviors^22,23,26–29,41^.

## RESULTS

### Convict cichlids can perform a goal-directed object manipulation task

We first tested the convict cichlids’ ability to solve a simple puzzle-box opening task (Figure 1, Supplementary Movie 1). Sexually mature individuals of both sexes were trained for multiple daily sessions of 5 trials of 5 minutes each (see “Methods” section). Briefly, the setup consisted of a box closed by a hinged lid (Figure 1a-e) that can be moved from side to side using a tab (Figure 1d, arrow). A food pellet was placed inside the box before the start of each trial. This goal-directed object-manipulation task requires relatively fine motor skills to manipulate the lid in a way that will allow the individual to access the food inside, and is analogous to previously used simple “puzzle-boxes” in the mammalian and avian literature^8,9,42^.

**Fig. 1.**
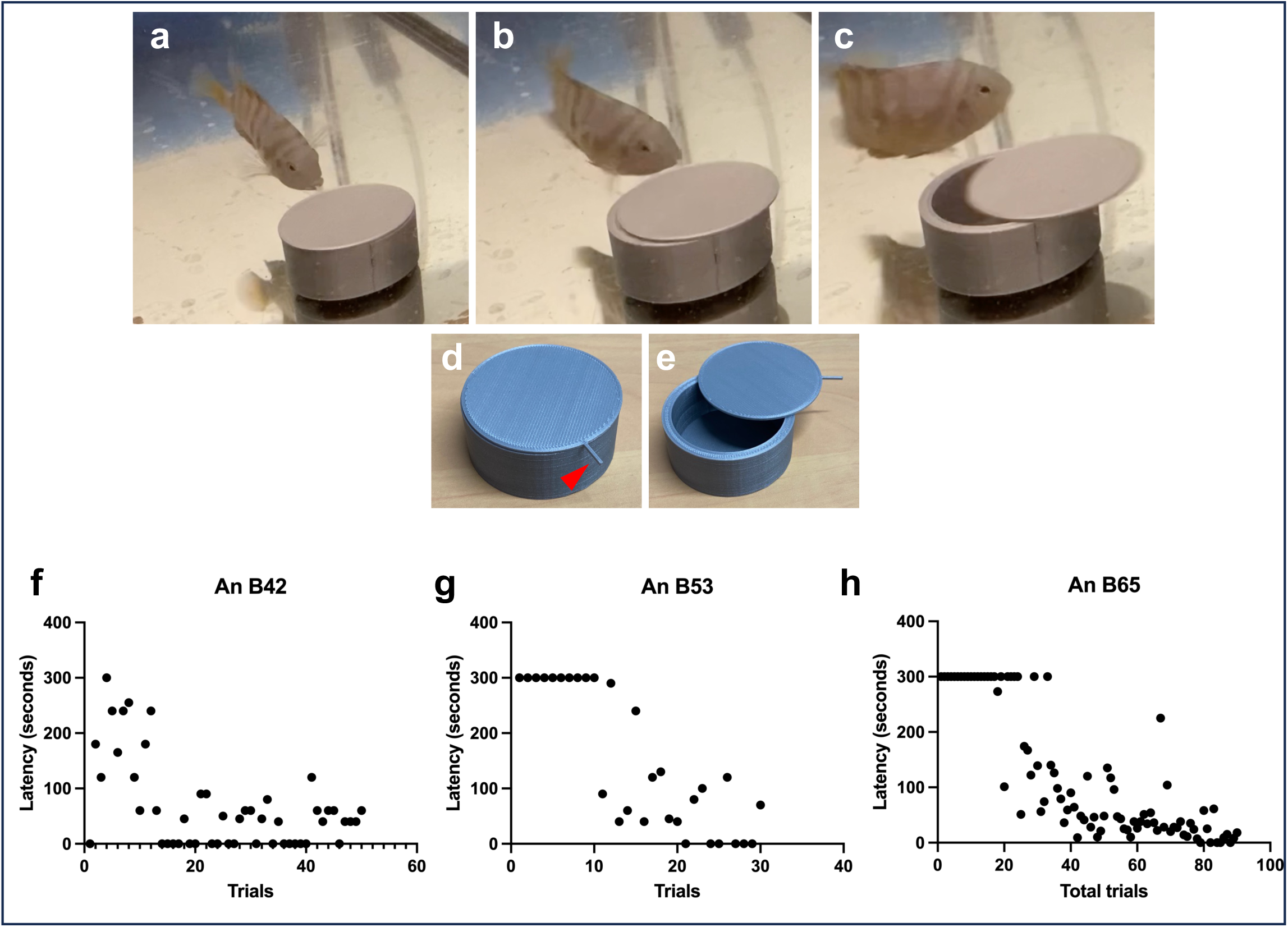
Goal-directed object manipulation task in the convict cichlid (*Amatitlania nigrofasciata*). (a-c) Sequence of motions used by convict cichlids to open the puzzle-box. Stills from Supplementary Movie 1. Cichlids first grabbed the lid’s tab (a), then performed a swinging motion to swing open the puzzle-box (b,c). (d,e) Puzzle-box used for the goal-directed object manipulation task in its closed (d), and opened (e) position. The tab (d, red arrow) on the lid can be used to open the box more easily. (f-h) Representative performance of cichlids during behavioral training. Latency to open the box is measured for each trial. Fish usually start by performing poorly (taking 5 minutes or more to open the puzzle-box), but show a rapid improvement until they are able open the puzzle-box in a few seconds (see Supplementary Figures 1-6).

All individuals (n=53) were able to open the box after being habituated to it for multiple hours in the first days of training. Latency to open the box was measured for each trial. In 44 individuals, representing 83% of all individuals included in this study, latency significantly decreased with training, going from 213 ± 116 seconds (n=880 trials) in the first 20 trials of training to 41 ± 64 seconds in the last trials of training (n=827), demonstrating that the convict cichlids had learned to open the box faster (Figure 1f-h, Wilcoxon signed rank test, p<0.05 in all cases, see Supplementary Figures 1-6). Latency during the final training sessions was measured on the last 20 trials for 39 individuals. For individuals who completed fewer than 40 total trials, latency was measured over the last 10 trials (n=4 individuals) or the last 7 trials (n=1 individual), depending on the number of trials completed. Anecdotally, we observed that the movements of fish became more efficient as training progressed. In the initial sessions, fish often struggled to open the box, whereas in later sessions they performed quick and accurate movements, resulting in much faster openings (see Supplementary Movies 1 & 2).

While all individuals were capable of opening the box given enough time, not all of them were able to learn to do so within the 5 minutes of each trial. Of the 53 individuals studied, 9 did not show any significant decrease in the latency to open the box, with 6 individuals displaying decreases in latency early during training but followed by irregular performance (see Supplementary Figures 1-2).

These results suggest that convict cichlids are capable of learning to perform a relatively simple problem-solving task requiring fine object manipulation, an ability that has seldom been studied in teleost fish.

### Increased neuronal activation in the lobar region of cichlids during a goal-directed object manipulation task

In order to identify the brain regions involved in this goal-directed object manipulation behavior, we used manganese-enhanced magnetic resonance imaging (MEMRI), which uses manganese to label activated neurons^43^. Following training (see previous section), n=35 convict cichlids were assigned to 3 different groups: an Experimental group (n=10) which would perform the problem-solving task, a Box control group (n=12) which would be exposed to the baited box without its lid, and a Food control group (n=13) which would be given the same amount of food as the Experimental group without the presence of the box (see “Methods” section). Both the Experimental and Food control group included n=3 fish each whose training performance was irregular, but who nonetheless performed the object manipulation behavior. The Food control group included an additional n=3 fish that were not able to complete the task within the 5 minutes trials but showed strong food motivation.

48 hours prior to imaging, the fish were injected with a solution of 50 mg/kg of MnCl_2_. Because MEMRI measures the accumulation of manganese, the fish were made to perform their assigned behavior as many times as possible to maximize manganese uptake by activated neurons. Specifically, the fish performed two behavioral sessions 24 hours before imaging and a final behavioral session one hour before imaging. At the end of this last session, they were anesthetized and kept in a Falcon 50 mL tube for the duration of the acquisition, which was done on an ultra-high field 17.2 T system^43^ (see “Methods” section).

After data normalization (see “Methods” section), we measured the relative average signal intensity of four regions of interest, in line with our previous study: the telencephalon, the optic tectum, the cerebellum, and the lobar region including the inferior lobe^29^. Normality was tested using the interquartile range method, and a search for outliers in the data led to the exclusion of two individuals in the Box control group and three individuals in the Food control group, resulting in a group size of n=10 for each of the three individual groups. Significant differences in relative signal intensity between groups were found using a two-way ANOVA (degrees of freedom: 2, f-value: 30.249, p<0.0001).

Significantly higher neuronal activation in all brain structures was found in the Experimental group compared to the Food control group (Figure 2, Tukey’s post hoc test, p<0.01 in all four structures), indicating that the increased activity in these brain regions in the Experimental group was due to other factor(s) than solely food intake, since both groups ate the same amount of food at the same rate.

**Fig. 2.**
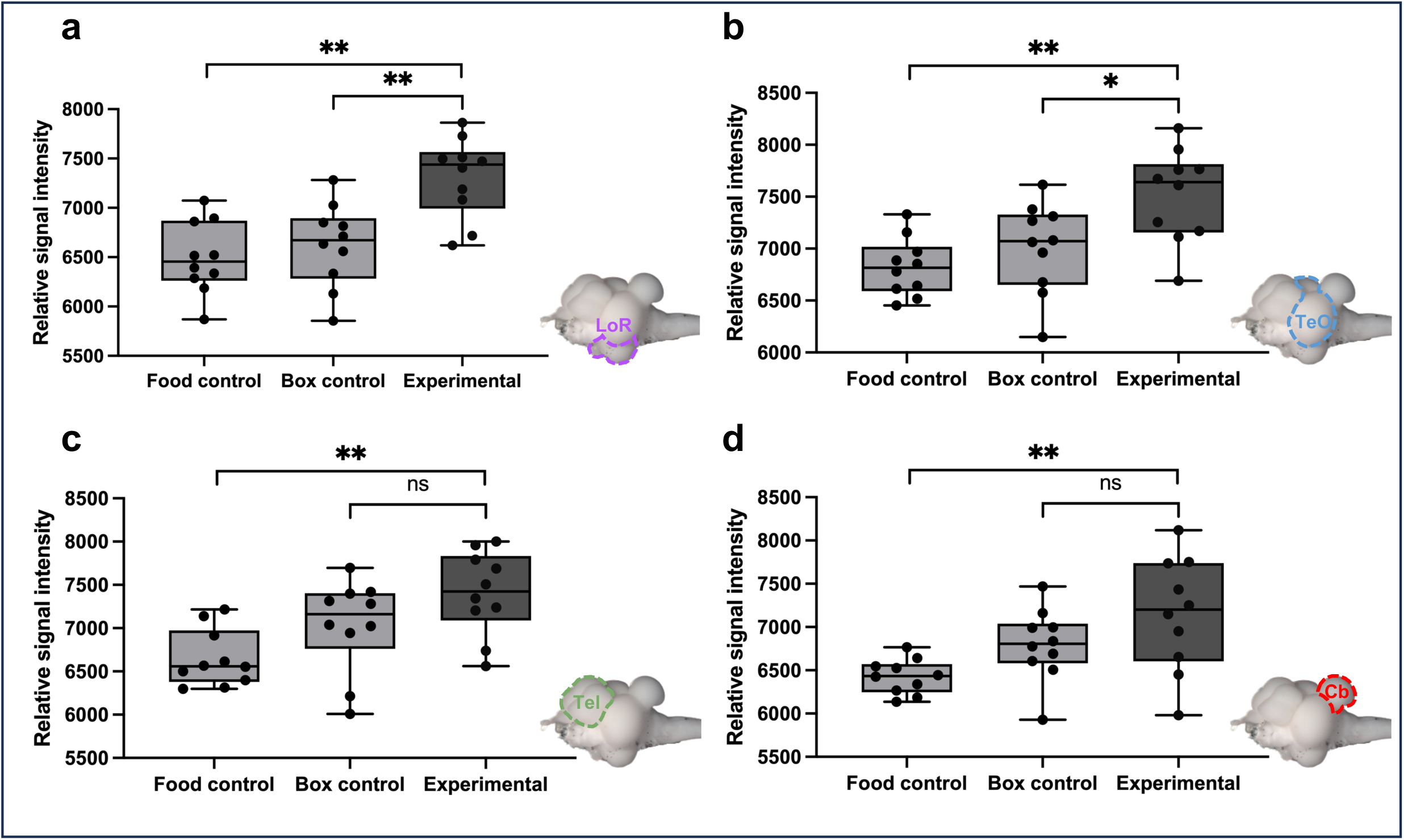
MEMRI reveals increased neuronal activation in the lobar region during a goal-directed object manipulation task. Box plots of the relative signal intensity measured for each group (n=10 in each group) in four brain regions: the lobar region (LoR) (a), the optic tectum (TeO) (b), the telencephalon (Tel), (c), and the cerebellum (Cb), (d), compared using two-way ANOVA and post-hoc Tukey’s test. Box plots indicate the minimum and maximum values, interquartile range (Q1 and Q3) and median value. (a) Significantly higher relative signal intensity in the lobar region of the Experimental group compared to both controls (p<0.01). (b) Significantly higher relative signal intensity in the optic tectum of the Experimental group compared to Box control (p<0.05) and Food control (p<0.01) groups. No significant difference in relative signal intensity in the telencephalon (c) and cerebellum (d) between Experimental and Box control groups (p=0.19 & p=0.22, respectively). *p<0.05, **p<0.01, ns: not significant

When comparing the Experimental group to the Box control group — where fish were given food inside the box without its lid — significantly higher neuronal activation was found in the lobar region, and to a lesser extent, in the optic tectum of the Experimental group (Figure 2a-b, lobar region: p<0.01, optic tectum: p<0.05). No significant difference in neuronal activation was found in the telencephalon or in the cerebellum between these two groups (Figure 2c-d, telencephalon: p=0.189, cerebellum: p=0.225). No significant differences were found when comparing the Box and Food control groups (Figure 2, p>0.05 in all cases).

Overall, by comparing different levels of the task: food intake (Food control group), eating from inside the box (Box control group), and opening the box (Experimental group), these results strongly suggest that the lobar region of cichlids is involved in this goal-directed object manipulation task, as it is significantly more activated in the Experimental group compared to both controls. This increase in activation is thus not due to food intake nor to the presence of the box itself, and more likely results from the action of opening the lid of the box (Figure 3).

**Fig. 3.**
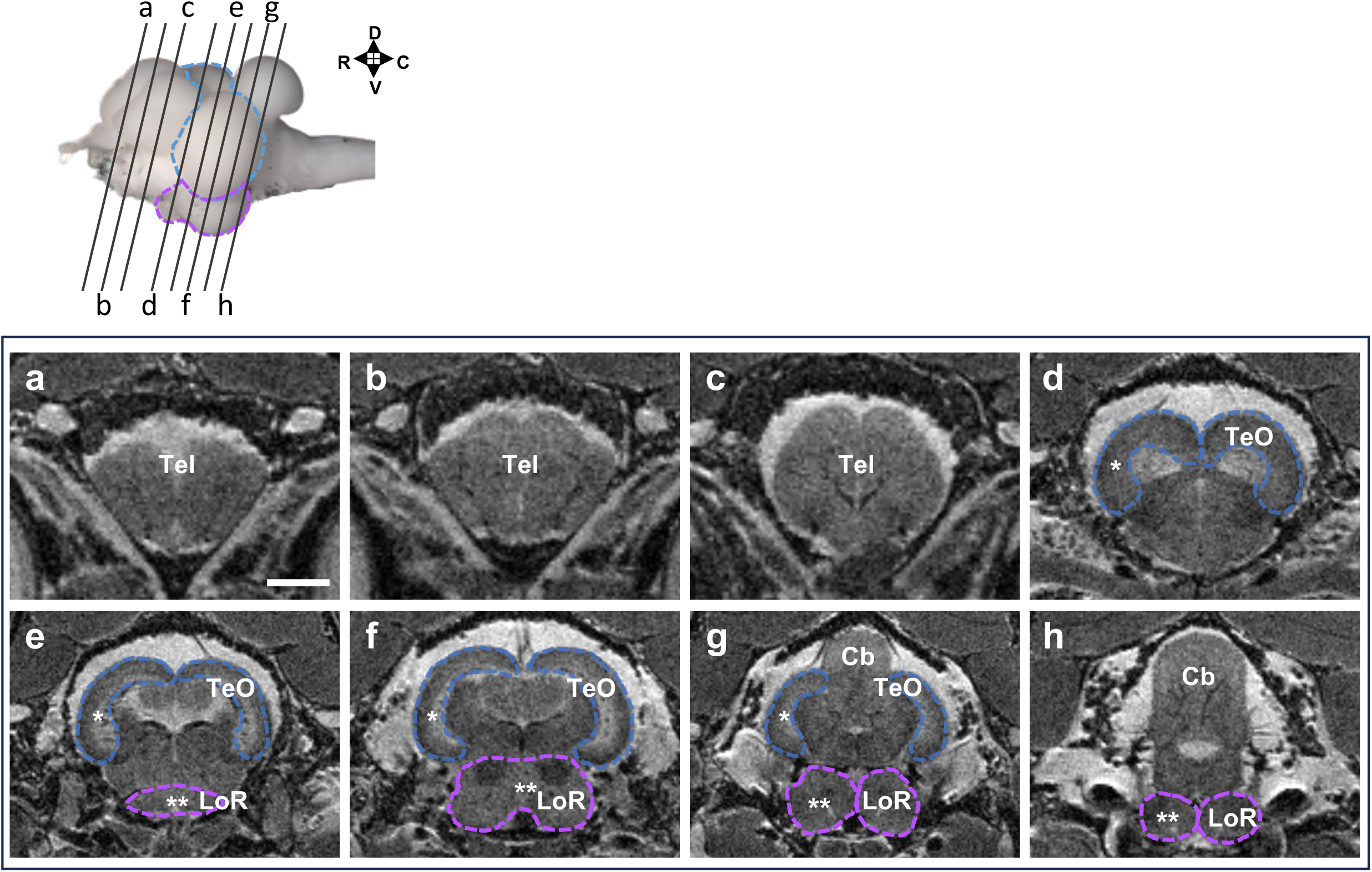
MRI neuroanatomical of the convict cichlid brain acquired at 17.2. The upper panel shows the approximate rostro-caudal position of each section, with the optic tectum (blue) and lobar region (purple) highlighted. (a-h) T2 weighted images acquired at 17.2 T in the axial plane. The optic tectum (TeO, blue), and lobar region (LoR, purple), which display increased neuronal activation in the behavioral task, are highlighted. Scale bar: 2.5 mm. Cb: cerebellum; LoR: lobar region; Tel: telencephalon; TeO: optic tectum. D: dorsal; V: ventral; R: rostral; C: caudal.

Along with the lobar region, the optic tectum was also significantly more activated in the Experimental group compared to the two controls. In teleost fish, the optic tectum is a large, laminated structure that receives multiple sensory modalities and is involved in visual processing to a much larger extent than its mammalian homolog, the superior colliculus^44^. This result possibly indicates that this goal-directed object manipulation task is heavily visually guided.

### Ultrasonic lesions of the inferior lobe support its role in goal-directed object manipulation tasks

To verify the role of the lobar region in the behavioral task, we used MR-guided HIFU to perform lesions of this structure^43^. This method was chosen over electrolytic surgical lesions as the latter required a craniotomy which convict cichlids do not tolerate well in our experience. MR-guided HIFU allows for precise lesions without skull opening, thus limiting potential pain for the animal and eliminating the risks of post-lesion infections, which are high in an aquatic environment (see “Methods” section).

A total of n=18 convict cichlids were selected for this experiment, divided into three groups. All fish were trained to perform the goal-directed object manipulation task (see “Methods” section) and showed a significant decrease through training in the time it took them to complete the task. We selected individuals which were reliably fast at performing the task, as our goal was to determine the impact of a lesion on their object manipulation abilities. Fish were anesthetized and placed in a specifically designed holder to expose the top of their head while keeping the rest of their body submerged, and an ultrasound transducer was placed on top of the fish’s head. Anatomical structures were identified and targeted using anatomical images acquired on a 7 T preclinical system (Bruker Biospin)^43^ (see “Methods” section). HIFU were applied for 15 seconds with an acoustic pressure of 6 MPa. MR-thermometry was used to assess the increase in temperature produced by the ultrasounds.

One group received bilateral lesions to the inferior lobe (Inferior lobe group, n=6, Figure 4a-f), which corresponds to the ventro-caudal part of the lobar region. A second group received lesions to the cerebellum, a structure that did not show increased activation in the MEMRI experiment (see previous section) to be used as control (Cerebellum group, n=6, Figure 4g-j). A third group, initially planned to receive bilateral lesions to the dorso-anterior part of the lobar region, was used as a Control group (n=6) because lesions were unsuccessful due to the small size of the targeted structure. These fish underwent the MR-guided procedure without lesions. Following the lesion, the fish recovered overnight and then underwent 4 sessions of 10 trials each of the behavioral task over 48 hours. When possible (n=16), following the last behavioral session, T2-weighted images of the animals were acquired, with T2 hypersignal being indicative of a lesion (Figure 4a-d, g-h), and the brain was sampled to visually confirm the presence and extent of the lesion (Figure 4e,i).

**Fig. 4.**
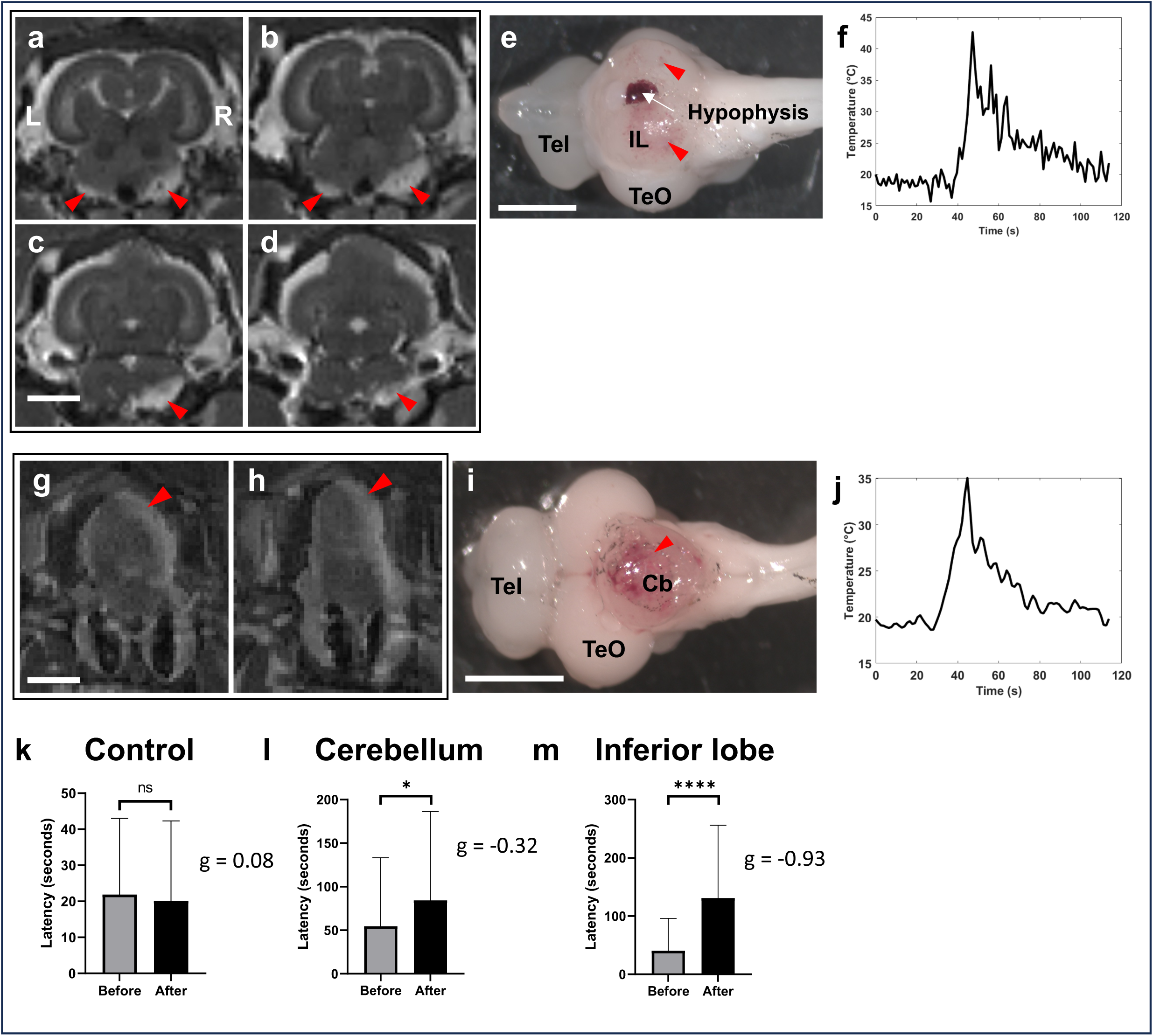
MR-guided High Intensity Focused Ultrasound (HIFU) lesions of the inferior lobe impair goal-directed object manipulation. Representative bilateral lesion of the inferior lobe (a-f), and of the cerebellum (g-j). (a-d) T2 weighted images acquired at 17.2 T 48 hours post-HIFU of an inferior lobe-lesioned cichlid showing bilateral hypersignal, consistent with HIFU lesion-related edema (red arrows). (e) Ventral view of the perfused brain of the same inferior lobe-lesioned convict cichlid as in (a-d), demonstrating extensive bleeding in the inferior lobe (red arrows). The white arrow indicates the hypophysis. (f) Temperature evolution in the inferior lobe of the individual in (a-e) over time during MR thermometry scan, showing an increase in temperature in the inferior lobe of around 22° C above baseline body temperature. (g-h) T2 weighted images at 7 T acquired 48 hours post-HIFU of a cerebellum-lesioned convict cichlid, showing hypersignal in the cerebellum (red arrows), consistent with the presence of HIFU lesion-related edema. (i) Dorsal view of the perfused brain of the same cerebellum-lesioned convict cichlid as in (h-i), demonstrating extensive bleeding in the cerebellum (red arrow). (j) Temperature evolution in the cerebellum of the individual in (h-j) over time during MR thermometry scan, showing an increase in temperature of around 15° C above baseline body temperature. (k) Before and after HIFU performance of the Control group, which underwent the procedure without brain lesions being obtained. No significant difference in latency was found when comparing the last 20 trials of training before HIFU lesion to the last 20 trials following the procedure in the control group. Wilcoxon rank sum test, n=6. The effect size (Hedges’ g) was likewise negligeable. (l) Before and after HIFU performance of the Cerebellum group. Comparing the last 20 trials of training before HIFU lesion to the last 20 trials following the procedure showed a modestly significant increase in latency required to open the puzzle-box following HIFU lesioning. Wilcoxon rank sum test, n=6. The effect size (Hedges’ g) was small (–0.32). (m) Increased latency in opening the puzzle-box in the goal-directed object manipulation task following inferior lobe lesions. The last 20 trials of training before the lesion were compared to the last 20 trials following HIFU lesions of the inferior lobe, with a significant increase in latency in the « After » condition. Wilcoxon rank sum test, p<0.0001, n=6. A large effect size was found by calculating Hedges’ g (–0.93). * p<0.05**** p<0.0001 g: Hedges’ g value measuring effect size. Scale bar: 2.5 mm. Cb: cerebellum; IL: inferior lobe; Tel: telencephalon; TeO: optic tectum. L: left; R: right.

We compared the last 20 trials of training to the last 20 trials following lesion, to account for the potential impact of the procedure and anesthesia itself in the first trials following lesion. Overall, all cichlids were capable of performing at least one trial in the behavioral task following lesions, albeit with large differences in the latency to complete the task between groups.

The Control group, having received no detectable brain lesion, performed at the same level as before the procedure (Figure 4k, mean latency 21.9 s ±21.2 s in the last 20 trials of training, 20.2 s ± 22.1 s in the last 20 trials following the procedure, Wilcoxon rank sum test, p=0.25). In the Cerebellum group, a slightly significant difference (Wilcoxon rank sum test, p=0.029) was found, with an increase in the latency required to open the box (Figure 4l, 54.5 s ± 78.7 s before lesion, 84.2 s ± 102 s following the lesion).

Animals in the Inferior lobe group were the most negatively impacted by the lesions, with a highly significant increase in the latency required to open the box (Figure 4m, animals went from 40.5 s ± 55.5 s before to 131 s ± 125 s following the lesion, Wilcoxon rank sum test, p<0.0001). Additionally, the effect size (Hedges’ g) was large in the inferior lobe group (g=-0.93) while it was small in the cerebellum group (g=-0.32), demonstrating that the impact on behavioral performance of inferior lobe lesions was much higher than cerebellum lesions.

While this was not quantified, we did observe that fish with lesions in the inferior lobe appeared less efficient in their box opening technique. Trained fish usually perform a singular swinging motion of the lid (Supplementary Movie 1), while inferior lobe lesioned individuals seemed to struggle to perform a smooth motion, instead needing multiple tries to succeed in opening the lid (Supplementary Movie 3). By contrast, cerebellum lesioned individuals appeared unaffected in the efficiency of their box opening movements (Supplementary Movie 4). Of note, no impact of the lesions was noted on locomotion, as the fish swam normally, nor on food motivation, as even the fish who performed poorly in the behavioral task readily ate food that was offered to them. Visual perception similarly seemed unaffected, as fish could navigate towards food and avoid a hand approaching them in their tank.

Although our initial design included a third lesion group targeting the dorso-anterior portion of the lobar region, technical limitations prevented successful lesioning in this area. Consequently, the current data do not allow us to fully assess the potential involvement of the entire lobar region in the task, which might not be limited to the inferior lobe.

Overall, this lesional experiment supports the MEMRI findings and point to the lobar region, and specifically to the inferior lobe, as playing a major role in goal-directed object manipulation tasks. Restricting the previous MEMRI analysis to the inferior lobe instead of the entire lobar region showed a highly significant increase in neuronal activation in the Experimental group compared to the Box control group that was given food inside the open puzzle-box (Figure 5, p<0.001). Notably, this increase in activation was even more significant than that of the lobar region as whole (Figure 2a).

**Fig. 5.**
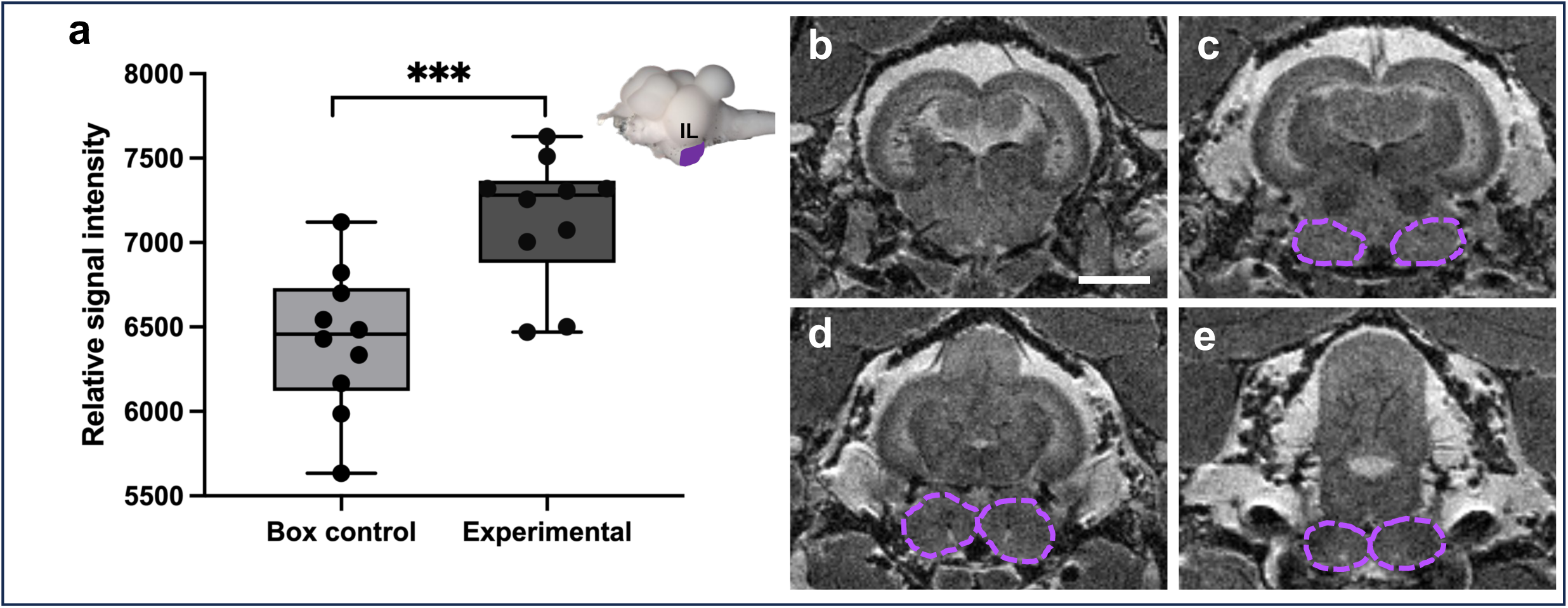
The caudo-ventral part of the inferior lobe is particularly activated following a goal-directed object manipulation task. (a) Highly significant increase in relative signal intensity in the caudo-ventral inferior lobe of the Experimental group compared to the Box control group (two-way ANOVA and post-hoc Tukey’s test, n=10 in all groups (Food control group not shown)). Box plots indicate the minimum and maximum values, interquartile range (Q1 and Q3) and median value. (b-e) T2-weighted images of the convict cichlid brain acquired at 17.2T with the caudo-ventral part of the inferior lobe highlighted in purple. Scale bar: 2.5 mm. *** p<0.001

## DISCUSSION

Object manipulation is a crucial step in evolving complex behaviors, including tool use. In mammals, object manipulation abilities are the most developed in primates, in which they have been the most studied^6,39,40,45^. Neuroanatomically, object manipulation in mammals is an overwhelmingly telencephalic behavior. In primates, the network for grasping and manipulating involves mostly cortical (premotor cortex and parietal cortex, especially the inferior parietal lobule and secondary somatosensory cortex)^39,40,46^ and subcortical telencephalic structures (putamen and globus pallidus)^47,48^, along with the cerebellum^49–52^.

In fish, object manipulation abilities are limited by the lack of prehensile limbs. The forelimbs of fish (i.e., pectoral fins) are adapted for swimming (or walking^53^). Although pectoral fins can be modified to a certain degree^54^,no fin has evolved into a prehensile appendage in teleosts. This leaves only the mouth to manipulate the environment with limited degrees of freedom^55,56^ compared to a mammalian hand^57^ or to the arm of an octopus^58^. Nevertheless, cichlids and wrasses can be relatively dexterous in manipulating objects with their mouths. The complexity of these behaviors seems to be correlated with the relative size of their inferior lobe: tool using wrasses have a larger inferior lobe than object manipulating cichlids, which in turn have a much larger inferior lobe than non-manipulative zebrafish^29^.

Our study reveals that the inferior lobe of convict cichlids, a non-telencephalic structure without homologs in mammals, plays a central role in goal-directed object manipulation. By combining MEMRI with targeted MR-guided HIFU lesions, we showed that the inferior lobe in cichlids can support behaviors associated with the telencephalon in amniotes.

In fact, the inferior lobe was long considered as part of the hypothalamus, with authors claiming a homology with the lateral hypothalamus of mammals^35,59^. With this assumption of homology, biting and snapping behaviors elicited by electrical stimulations of the inferior lobe were interpreted as food intake behaviors^60,61^. However, a developmental study conclusively showed that only the inner-most cell populations of the inferior lobe along the hypothalamic ventricle are of a hypothalamic origin, while most cells in the external part of the inferior lobe are in fact of a mesencephalic origin^21^. It is this external part of the inferior lobe that becomes enlarged in cichlids and wrasses^21,29^. Thus, the inferior lobe is mostly mesencephalic instead of hypothalamic, and has no homolog in tetrapods. These developmental findings called into question the functional role of the inferior lobe.

In our study, the increased neuronal activity in the inferior lobe was specifically associated with the performance of the object manipulation task, rather than with food intake. Fish exposed only to food in the same quantities as the Experimental group (Food control group) did not show comparable levels of activation, indicating that the act of opening the box — a coordinated, goal-directed behavior — drove inferior lobe activation. Likewise, lesions of the inferior lobe did not reduce food motivation in convict cichlids. Intact food motivation thus tends to disprove the putative role of the inferior lobe as a food motivation center^35,59–61^.

Based on our HIFU lesion results, the inferior lobe didn’t appear to be functionally related to locomotion either, as lesioned animals did not show discernible deficits in swimming behaviors. Similarly, visual perception was unaffected in inferior lobe-lesioned animals, who were able to navigate towards and catch food pellets dropped into their tanks, or avoid a hand approaching them. In the case of cerebellum lesions, the lack of effect on locomotion, while surprising when compared to similar lesions in amniotes which produce extensive motor deficits^62,63^, is consistent with previous lesion studies in goldfish and trouts^64–66^. In these, the corpus cerebelli, the region of the cerebellum that we lesioned in the present study, was surgically ablated, with no significant impact on swimming behavior in lesioned animals.

As lesions to the inferior lobe did not seem to affect food motivation, general locomotion, and gross visual perception, the observed deficits in the goal-directed object manipulation task were not due to widespread impairments in movement or simple sensory processing. They instead point to specific disruptions of the circuits supporting higher-order, sensory motor integration. This is in line with connectivity findings pointing to the inferior lobe as a putative multisensory integration center^30,31,34,67^ receiving visual^31,33–35^, gustatory^30,34,36,37^ and somatosensory^31^ information. This multisensory connectivity converges to the inferior lobe but also to the broader lobar region. Our previous study has showed that the inferior lobe and the lobar region were significantly enlarged in wrasses and cichlids compared to other teleosts like the zebrafish, and abundantly connected with the pallium^29^. This suggests that in wrasses and cichlids, who display multiple examples of higher-order cognition, including tool use in wrasses, the lobar region might be involved in more complex behavioral functions than in zebrafish, such as goal-directed object manipulation tasks.

We initially focused our MEMRI analysis on the broader lobar region, but lesions targeting the inferior lobe rather than the entire lobar region pointed to a specific role of the inferior lobe. Further MEMRI analysis of the inferior lobe itself showed that activation was even more significantly elevated in the inferior lobe during the goal-directed object manipulation task compared to the lobar region as a whole. Lesions of the inferior lobe mostly ablated the nucleus diffusus (nucleus diffusus lobi inferioris, NDLI^68^), the most caudal part of the lobar region. The NDLI projects heavily, although indirectly, to the cerebellum via the nucleus centralis lobi inferioris pars anterior (NCa) and the lateral valvular nucleus^68^, which can explain the slight effect of cerebellar lesions. However, while cerebellum-lesioned fish also showed a mild deficit in the task, the magnitude of the impairment was substantially greater in the inferior lobe group (Supplementary Movies 3-4). Behavioral observations further revealed qualitative differences: whereas cerebellum-lesioned fish typically executed a smooth, singular movement to swing open the lid, inferior lobe-lesioned fish showed less efficient, repeated attempts, indicating a loss of fine motor coordination. The inferior lobe thus potentially allows for the integration of sensory cues with precise motor outputs necessary for delicate object manipulation.

We here described the puzzle-box opening task as “goal-directed object manipulation”. However, equivalent behavioral tasks were often considered “problem-solving tasks” in previous publications^8,9,42^. Thus, while we are careful in our interpretation, it is possible to consider this task as a form of simple problem-solving. Adding steps to increase the complexity of the puzzle-box would more clearly classify it as problem-solving.

Overall, the involvement of the inferior lobe in goal-directed object manipulation behaviors is particularly striking given its non-telencephalic origin with no homolog in tetrapods. This contrasts with data in amniotes where higher-order behaviors such as tool-use, problem-solving, and complex motor planning are functions exclusively mediated by telencephalic circuits, especially the pallium^3,39,52,69,70^. Our findings suggest that teleosts have evolved a different neural architecture to support analogous behavioral outcomes, recruiting non-homologous brain regions such as the inferior lobe. This supports the broader concept of evolutionary convergence at the level of behavioral functions, even when the underlying neural substrates differ substantially across vertebrate clades.

In conclusion, our study provides the first experimental evidence that the inferior lobe is involved in object manipulation in a goal-directed manner in teleost fish. These findings challenge the notion that such functions necessarily arise from the telencephalon, and broaden our understanding of how complex behaviors can emerge from distinct neural architectures across vertebrate evolution. Future investigations should aim to dissect the microcircuitry within the inferior lobe, determine its role in more complex forms of problem-solving, and explore how these findings might inform our understanding of cognitive evolution in other non-mammalian lineages.

## METHODS

### Study animals and housing

Sexually mature convict cichlid (*Amatitlania* nigrofasciata) of both sexes weighing between 2 and 17.2 grams (average: 6.72 ± 3.72 g) were obtained from commercial suppliers (Aquariofil.com, Nîmes, France). They were housed in large 120 to 400L filtered and aerated fresh water communal tanks, with water kept at 25° C. A light/dark cycle of 12/12 hours was maintained throughout the duration of the experiments. A week prior to the start of the behavioral tests, individuals were isolated in 40 L tanks placed next to each other so that visual contact could be maintained between individuals. Once the experimental protocol started, fish were only fed during the behavioral experiments. All procedures were conducted in compliance with the official regulatory standards of the French Government and the committees (CEA, NeuroPSI) under protocol number APAFIS #51934-2024111418122817 v5; APAFIS #55234-2025051210564718 v1 & APAFIS #40422-2023012000331085 v7.

### Behavioral training

During the box-opening training, only one behavioral session consisting of 5 trials was performed in one day. Training was performed starting one month before experiments, but the total number of sessions varied between individuals (due to how fast the fish would learn to open the box, but also MRI scheduling and experimenter availability). The minimum number of training sessions was 6 (total: 30 trials), while the maximum was 28 (total: 140 trials). Opaque separations were placed between tanks five minutes before session start so that fish could not see each other for the duration of the session. A food pellet was placed in a circular grey plastic 3D printed box (diameter: 4.8 cm, height: 2 cm) closed by a hinged lid that could slide open with a tab to facilitate manipulation (Figure 1d-e, Supplementary Movies 1-2). The box was placed in the fish tank. Fish were allowed 5 minutes to open the box, and time to open the box was recorded (see Supplementary Figures 1-6). Failure to do so was recorded as a timeout (included in the analysis) and the experimenter moved on to the next trial. At the end of each session, the opaque separations and box were removed. As fish were usually hesitant to approach the box in the first few sessions (5-10), the box was left at the end of the last trial inside the tank in case the fish didn’t open it, to habituate the fish to its presence. In almost all cases, after a few hours the fish had opened the box. All fish included in this study were trained to open the box, and all the individuals tested could successfully do so. In all but a few cases, training resulted in the latency to open the box decreasing significantly. This was tested using a Wilcoxon rank sum test (a shapiro-Wilk test determined that data did not follow a normal distribution) comparing the average latency of the first 20 trials to the last 20 trials of training. In some cases, when fish had fewer than 40 trials, the first 20 trials were compared to the last 10 trials (n=4 individuals) and in one individual, to the last 7 trials. All Tests were performed using R Studio v.2023.12.1.

### Manganese-Enhanced MRI (MEMRI) behavioral procedure

After training, an Experimental group, along with two control groups were formed (n=10 in the Experimental, n=13 and n=12 in the Food and Box control groups, respectively).

In the Experimental group, a behavioral session consisted of 10 trials of 5 minutes each. Fish were rewarded with food in only half the trials to avoid satiety. The closed box was placed in the tank, latency to open the box measured, and trials ended after 5 minutes, regardless of whether the box was opened or not. As fish usually had opened the box within 2 minutes of each trial, some additional manipulation of the box was frequently observed, with the fish biting the lid and moving it from side to side or detaching it from the box completely.

In the Box control group, the box without the lid was used to dispense food and therefore getting the food pellet did not require any manipulation. This group was intended to distinguish neuronal activity related to the presence of the box and to eating from inside the box from neuronal activity related to the box opening behavior in the Experimental group. A behavioral session consisted of 5 trials of 10 minutes each, all including food distribution. Food was placed at the bottom of the box, and the box was placed in the tank. As soon as the fish ate the food pellet, the box was removed from the tank to avoid manipulation.

In the Food control group, the food pellet was dropped inside the tank without the box. This group was intended as a baseline, to distinguish neuronal activity related to food intake from the activity related to the box opening behavior in the Experimental group. A behavioral session consisted of 5 trials of 10 minutes each, with each trial starting as soon as the food hit the water.

### MEMRI acquisitions

In order to visualize neuronal activity related to the box opening behavior, manganese-enhanced magnetic resonance imaging (MEMRI), was used^71^. MEMRI uses manganese, a magnetic resonance contrast agent and calcium analog, to label active neurons.

48 hours prior to imaging, fish were anesthetized in MS222 (170 mg/L), weighed, and injected intraperitoneally with a solution of 100 mM MnCl_2_ in 1 M HEPES (pH 7.5, 6 µL/g: 50 mg/kg of MnCl_2_). Fish were given 24 hours to recover, then two behavioral sessions were performed 24 hours prior to imaging, spaced approximately 6 hours apart. On the day of imaging, a final behavioral session was performed one hour before imaging. At the end of the session, the animals were anesthetized in MS222 (180 mg/L), then transferred in a 50 mL Falcon tube filed with fresh anesthesia water. Depending on the size of the animal, a paper towel was wrapped around the fish to prevent motion inside the tube.

MR acquisitions were performed on a 17.2 T Bruker imaging system (Bruker Biospin) equipped with an Avance III console running Paravision 6.0.1 using a 45-mm diameter birdcage transmit/receive volume coil (Rapid Biomedical). Two consecutive T1 weighted images were acquired with the following parameters: TE/TR = 2.8/250 ms, flip angle = 90°, 24 slices, slice thickness= 0.2 mm, in plane resolution = 0.08 mm x 0.08 mm, acquisition time = 26 min.

At the end of the acquisition, fish were ventilated with fresh water until voluntary movement started again, then transferred back into their housing tanks. The procedure from beginning of anesthesia to end of acquisition lasted approximately two hours.

### MEMRI data processing

The acquired data was processed using the FSL suite of tools^72^. The two T1 weighted images were averaged to produce a single, higher signal to noise image. To normalize the signal intensity across individuals, the mean intensity of the retina of each individual was measured by drawing a region of interest (ROI) on one slice in the middle of the eye, as the retina was reliably labeled with manganese in all fish. An arbitrary normalized intensity value of 10,000 was divided by the mean intensity of the retina, then the MRI data was multiplied by this value to obtain normalized images. FSLeyes was used to draw ROIs to compare manganese labelling in different brain regions between groups. The brain was manually segmented into five ROIs: the telencephalon, the optic tectum, the lobar region, the inferior lobe, and the cerebellum. The ventral end of the optic tectum was used as the dorsal boundary of the lobar region. The cerebellum was underestimated as its caudal end was not imaged, and we excluded the valvula cerebelli as it was not possible to distinguish it from the torus longitudinalis and other midbrain structures with the spatial resolution of our images. Structure boundaries were validated by a researcher blind to which group each individual fish belonged to. Mean intensity was measured for each ROI. Statistical tests (Two-way ANOVA and Tukey’s post-hoc test) were performed using RStudio v.2023.12.1+402.

### Magnetic resonance (MR)-guided High Intensity Focused Ultrasounds (HIFU)

After training, n=18 convict cichlids were separated into 3 groups: a Control group (n=6), a Cerebellum group (n=6) and an Inferior lobe group (n=6). Fish were anesthetized with MS222 (200 mg/L) and placed in a Falcon 50 mL tube filled with water with its top open, allowing for the top of the fish head to be out of the water while the gills remained submerged. The tube was placed inside a specifically designed, in-house built RF coil. An 8-element focused ultrasound transducer with a central frequency of 1.5 MHz, a geometrical depth of 20 mm and a diameter of 30 mm was placed atop the fish head, using ultrasound gel to seal any air gap. Additional details can be found in Estienne et al.^43^.

T2 weighted images were acquired with a 7 T Biospec MRI system (Bruker Biospin) with the following parameters: 3000 ms TR, 20.75 ms effective TE, RARE factor = 8, 0.25 x 0.25 mm^2^ in-plane resolution, 32 x 32 mm^2^ field of view, 19 axial slices with a thickness of 1 mm, total scan time of 3 min 12 s. Using these anatomical images, we targeted the cerebellum (Cerebellum group) or the inferior lobe (Inferior lobe group) by moving the transducer using a dedicated software (Thermoguide®, Image Guided Therapy, Pessac, France)^73^ and acquiring new T2 images to control the position of the transducer. In the Control group, we initially wanted to target the anterior part of the lobar region, but due to its small size, the ultrasonic shots missed their targets in all our 6 individuals. As a result, these individuals did not receive a lesion (verified by T2 weighted images 48 hours after the procedure) and were considered controls for the procedure itself.

Once targeting was confirmed, we launched an MR thermometry acquisition lasting 1 min 55 s consisting of a standard Fast Low Angle Shot (FLASH) sequence repeated every 1.28 s. The acquisition parameters for the FLASH sequence were as follows: TE 3.5 ms, TR 10 ms, flip angle of 30°, spatial resolution of 0.5 x 0.5 x 3 mm^3^, for a field of view 32 x 32 mm^2^. One single slice was acquired in the axial orientation. 30 seconds into this acquisition, the sonication process was launched with 15 800 ms shots separated by 200 ms, with an acoustic pressure of 6 MPa. Changes in signal intensity indicative of a rise in temperature were measured using Paravision software (PV6.01, Bruker Biospin, Germany) to confirm that lesioning indeed took place. As the inferior lobe is a bilateral structure, two series of shots were performed, one for each side. The Control group also received two series of shots.

After lesioning, the animals were awakened from anesthesia by ventilation with fresh water and placed in their home tanks with methylene blue to prevent infection, as sonication sometimes produced smalls burns on the skin directly in contact with the ultrasound transducer.

### Behavioral and anatomical assessments of HIFU lesions

Starting 24 hours after lesioning and up to 48 hours after lesioning, convict cichlids performed between 20 and 40 trials spread over 2 to 4 behavioral sessions (average: 36.7 ± 5.9 trials). Latency to open the closed puzzle-box was measured. The mean average latency of the last 20 trials following lesioning for each group was compared to the mean average latency of the last 20 trials of training. Normality was tested using a Shapiro-Wilk test, and as normality could not be verified, a Wilcoxon rank sum test was used to compare the performances before and after lesion in each group. Additionally, the effect size was measured for each group, to assess the strength of the lesion’s impact on the before and after behavioral performance. Given the small sample size (n=6 in each group), we used Hedges’ g as our effect size measure.

48 hours after HIFU lesions, the anatomical site of the lesion was checked by performing high-resolution T2-weighted RARE acquisitions using either a 7 T Biospec system or a 17.2 T system, when possible, and/or by dissecting the brain and visually checking for signs of hemorrhage. On the 7 T Biospec system, the following acquisition parameters were used: TE = 8 ms, TR = 1500 ms, field of view = 28 x 28 mm^2^, spatial resolution = 0.2 x 0.2 x 0.2 mm^3^, 15 axial slices.. On the 17.2 T system, the following acquisition parameters were used: TE = 6.84 ms, TR=3000 ms, field of view = 28 x 28 mm^2^, spatial resolution = 0.1 x 0.1 x 0.1 mm^3^, 18 axial slices. Lesions reliably produced T2 hypersignal. After acquisition, or if acquisition was not possible due to scheduling conflicts with the MR scanners, the animals were perfused transcardially with 10% formalin, the brain dissected and photographed under a dissection microscope.

### Statistics and reproducibility

All statistical analyses were performed using R Studio software v.2023.12.1+402. MEMRI data was also analyzed by a researcher blinded to the different groups. Graphs were created using R Studio, GraphPad Prism v.10.4.2 (GraphPad Software), and FSLEyes 1.14.0.

## DATA AVAILABILITY

Data reported in this paper will be shared by the corresponding author upon request. This paper does not report original code.

## ACKNOWLEDGEMENTS

We thank Anthony Novell, Corentin Cornu and the BioMaps laboratory (Paris-Saclay University, CEA, CNRS, Inserm, BioMaps, Service Hospitalier Frédéric Joliot, Orsay, France) for their help in calibrating the ultrasound transducer. We thank Dorian Champelovier for his help with designing and 3D printing the puzzle boxes used in this study, and Florian Razy-Krajka for critical reading of the manuscript. We also thank the PSI-CO platform (Cynthia Froc) at NeuroPSI for help with behavioral training, and the DECA team members.

This study was supported by INSB/CNRS Call “Diversity of Biological Mechanisms” and ANR EVONECTOME (ANR-23-CE37-0021).

## AUTHOR CONTRIBUTIONS

Conceptualization, K.Y. and P.E.; methodology, L.C., G.Pa. B.L., P.E., K.Y.; funding acquisition and supervision, K.Y. and L.C.; validation and visualization, K.Y., P.E., L.C., G.Pa., G.Po; fish behavioral training, P.E., T.D., A.D.; first draft of manuscript, P.E., and K.Y.; all authors contributed to data analysis, interpretation and revision of the manuscript.

## MATERIALS & CORRESPONDENCE

Correspondence and requests should be addressed to Kei Yamamoto.

