## Supplementary Figures for "Problem-solving without a cortex: inferior lobe drives goal-directed object manipulation in cichlid fish"

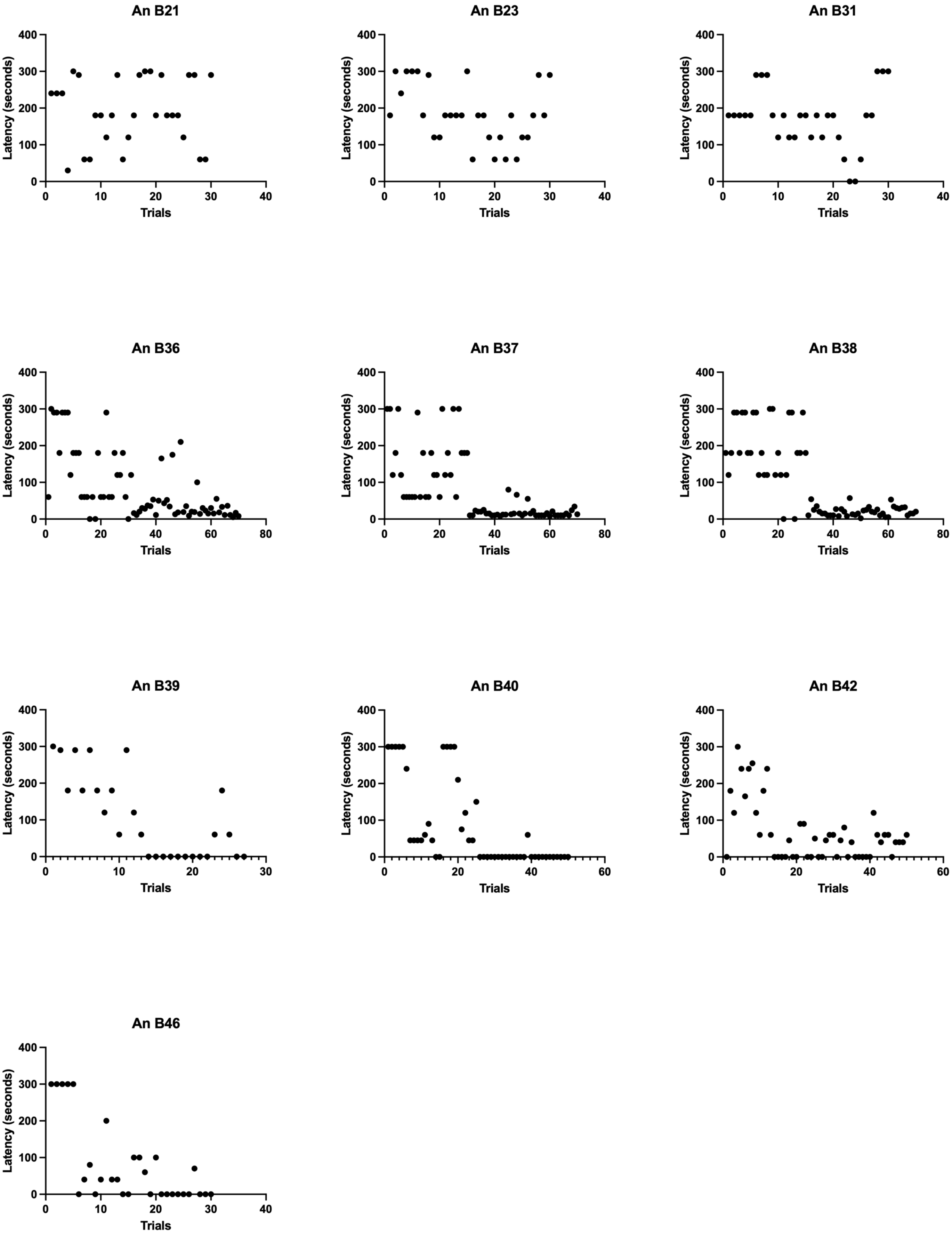


**Supplementary Fig. 1. Training data for all individuals in the MEMRI Experimental group.**

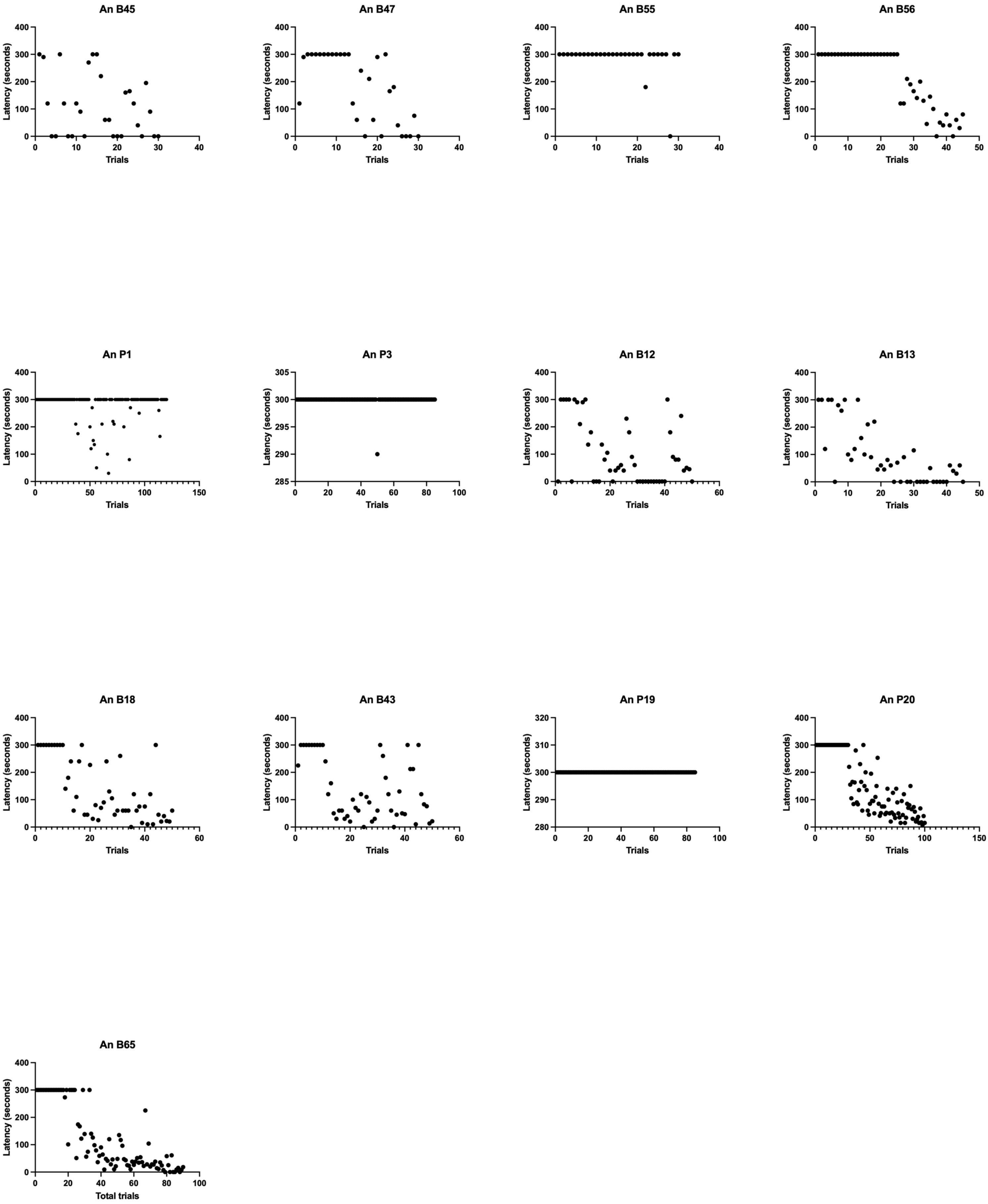


**Supplementary Fig. 2. Training data for all individuals in the MEMRI Food control group.**

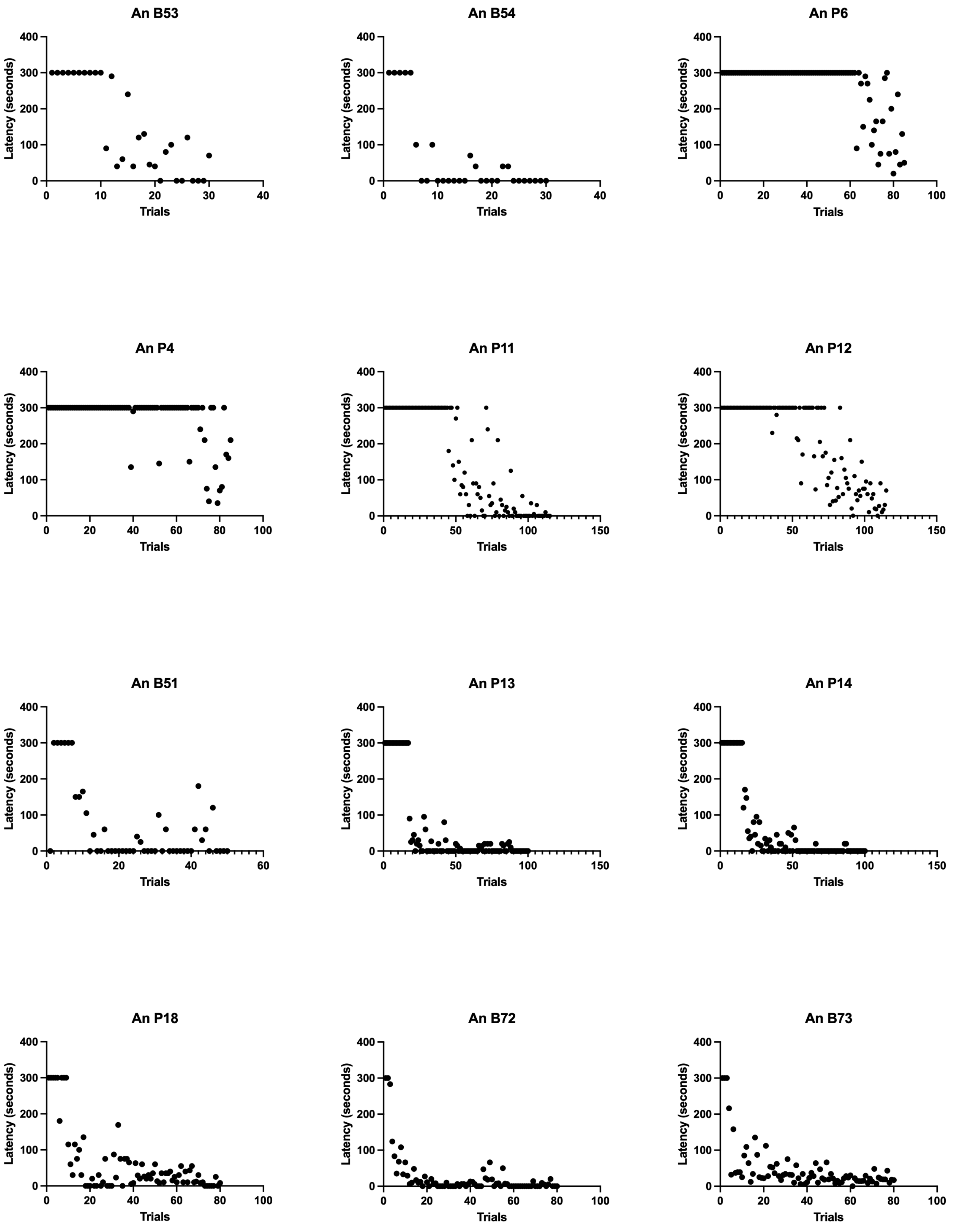


**Supplementary Fig. 3. Training data for all individuals in the MEMRI Box control group.**

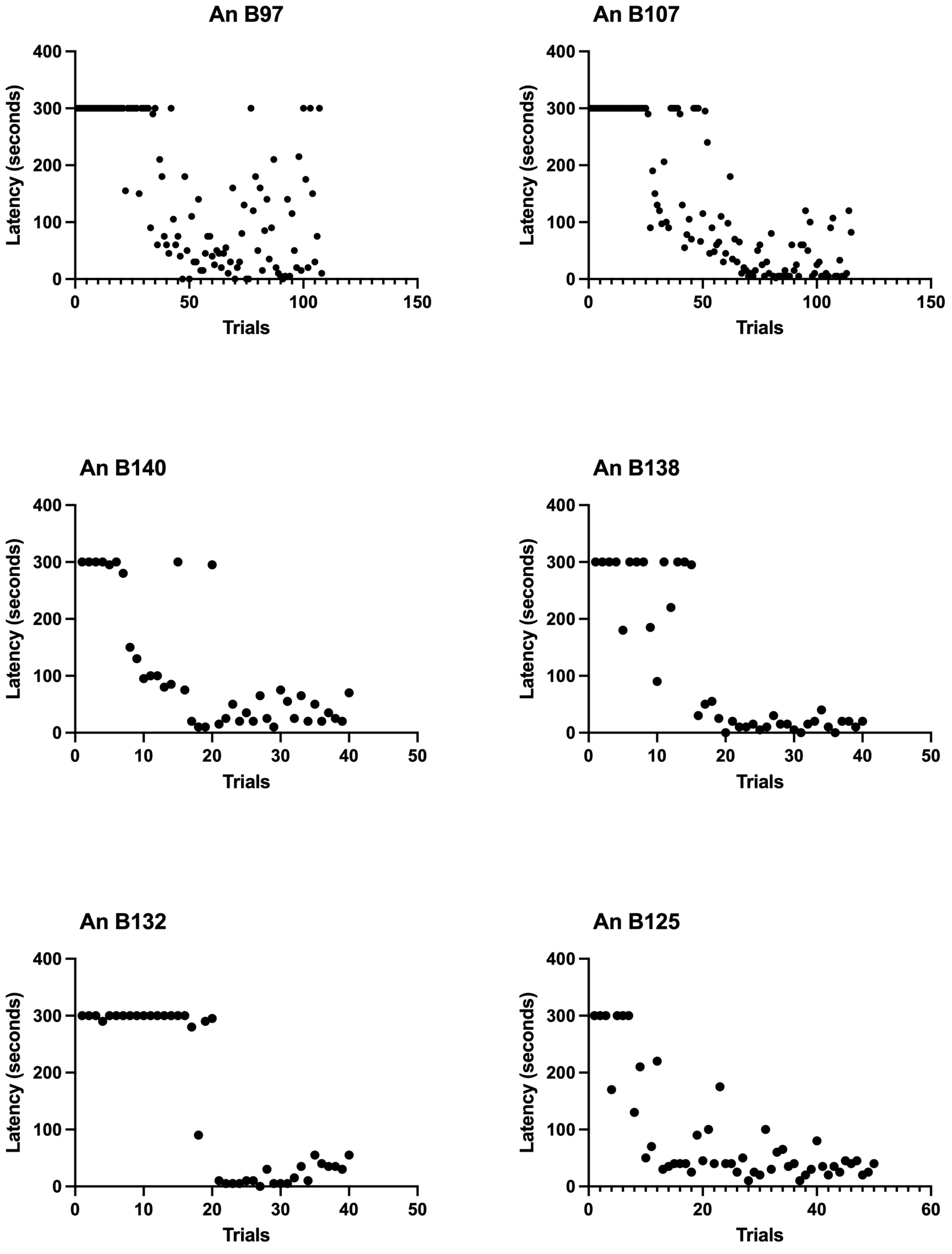


**Supplementary Fig. 4. Training data for all individuals in the HIFU Inferior lobe group.**

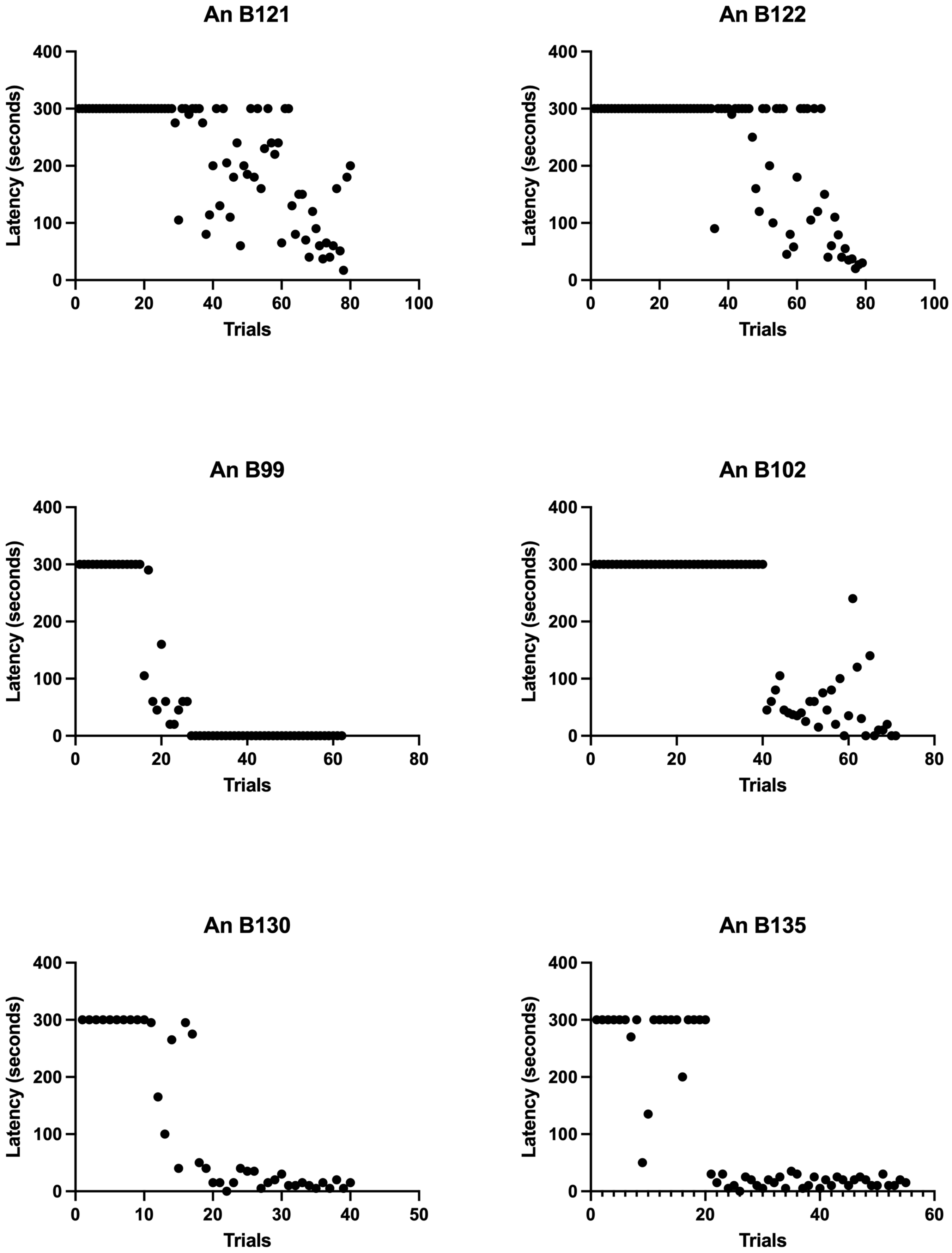


**Supplementary Fig. 5. Training data for all individuals in the HIFU Cerebellum group.**

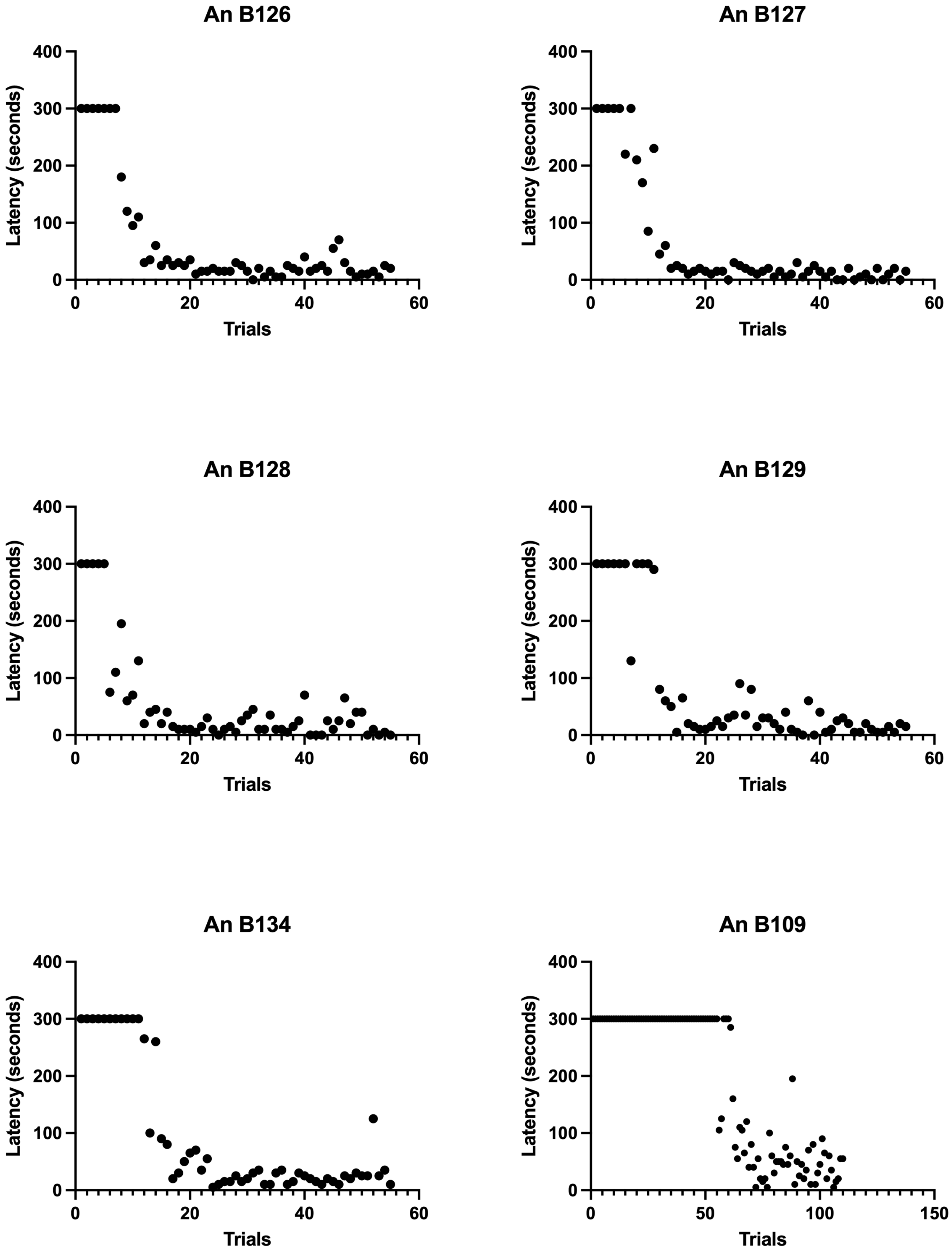


**Supplementary Fig. 6. Training data for all individuals in the HIFU Control group.**
